# Biofilms preserve the transmissibility of a multi-drug resistance plasmid

**DOI:** 10.1101/2022.04.18.488688

**Authors:** Genevieve A. Metzger, Benjamin J. Ridenhour, Michael France, Karol Gliniewicz, Jack Millstein, Matthew L. Settles, Larry J. Forney, Thibault Stalder, Eva M. Top

**Author notes:** <u>Corresponding Author</u>: Department of Biological Sciences, Life Sciences South 258, University of Idaho, Moscow, ID 83844-3051.

## Abstract

Self-transmissible multidrug resistance (MDR) plasmids are a major health concern because they can spread antibiotic resistance to pathogens. Even though most pathogens form biofilms, little is known about how MDR plasmids persist and evolve in biofilms. We hypothesize that (i) biofilms act as refugia of MDR plasmids by retaining them in the absence of antibiotics longer than well-mixed planktonic populations, and that (ii) the evolutionary trajectories that account for the improvement of plasmid persistence over time differ between biofilms and planktonic populations. In this study, we evolved *Acinetobacter baumannii* with an MDR plasmid in biofilm and planktonic populations with and without antibiotic selection. In the absence of selection biofilm populations were better able to maintain the MDR plasmid than planktonic populations. In planktonic populations plasmid persistence improved rapidly but was accompanied by a loss of genes required for the horizonal transfer of plasmids. In contrast, in biofilms most plasmids retained their transfer genes, but on average plasmid persistence improved less over time. Our results showed that biofilms can act as refugia of MDR plasmids and favor the horizontal mode of plasmid transfer, which has important implications for the spread of MDR.

## INTRODUCTION

Antibiotic resistance is widely recognized as one of the most serious problems facing healthcare today. Resistance to antibiotics, including those of last resort, often results from the acquisition of resistance genes by horizontal gene transfer (HGT) ^1,2^. When present as part of mobile genetic elements such as plasmids, resistance genes can rapidly spread to various species or strains of bacteria ^3,4^. This makes multidrug resistance (MDR) plasmids, particularly those with a broad host range, a major problem in fighting the spread of antibiotic resistance.

Understanding how these plasmids evolve in the presence and absence of antibiotics is critically important. In the presence of antibiotics, MDR plasmids are maintained in bacterial populations via positive selection ^5^, but when antibiotics are removed the fitness cost of plasmid carriage ^6–8^ is no longer offset by the benefit of antibiotic resistance ^9^. Therefore, plasmids are expected to be lost from bacterial populations via purifying selection unless (i) partitioning and post-segregational mechanisms limit the formation of plasmid-free cells, (ii) reacquisition of plasmids counteracts plasmid loss and cost, (iii) co-residing plasmids and other bacteria in the community can prevent plasmid loss ^10,11^, or (iv) the cost of plasmid carriage is reduced, eliminated or reversed by evolution ^12^. The rate at which plasmids are lost from a population probably depends on a combination of these factors and is inversely proportional to ‘plasmid persistence’ – the ability of a plasmid to maintain itself in the absence of known positive selection for the plasmid. Previous studies from our and other research groups have repeatedly shown that plasmid persistence can improve during growth in the presence or absence of antibiotics (for example, ^13–24^). However, almost all these studies used planktonic populations. Much less is known about how plasmid persistence evolves in spatially structured populations such as biofilms, even though they represent the most common form of bacterial growth and are the causes of many recalcitrant infections ^25–28^.

Due to spatial structure, bacteria that grow in biofilms only compete locally with their neighbors, which protracts selective sweeps and leads to increased genotypic and phenotypic variation within a population ^29–39^. Additionally, the gradients of nutrients and electron acceptors cause habitat heterogeneity that results in local adaptation of subpopulations. These biofilm features allow bacteria to access more fitness peaks in rugged adaptive landscapes. This can result in evolutionary outcomes that may not be observed in planktonic populations that routinely experience strong selective sweeps ^40–42^. Previously we showed that under antibiotic selection, evolution in biofilms results in a higher diversity of plasmid persistence phenotypes as compared to planktonic populations ^43^. In a separate study we have found that biofilms of plasmid-bearing cells maintain a greater diversity of mutations than planktonic populations. Among these diverse biofilm genotypes were clones that better retained their plasmid than any clone evolved in planktonic populations ^44^. These seminal studies showed how biofilm growth can affect the evolution of plasmids persistence. Because many bacterial pathogens with antibiotic resistance plasmids ^45,46^ form biofilms, we need to study the evolution of their plasmids under these growth conditions.

The biofilm-forming bacterium *Acinetobacter baumannii* is an emerging threat in the United States and worldwide because it causes wound infections, ventilator-associated pneumonia, and sepsis ^47,48^. Further, it is known to rapidly acquire new forms of antibiotic resistance by HGT ^49–51^ and pan-drug resistant strains have been documented ^52^. Despite the medical importance of *A. baumannii* and the role of HGT in its acquisition of antibiotics resistance, little information is available on how this organism and recently acquired MDR plasmids co-evolve.

Previously we found that the persistence of a broad-host range MDR plasmids pB10 in *A. baumannii* increased over time in both biofilm and planktonic populations treated with antibiotics, but this phenotype showed more diversity after evolution in biofilms ^43^. Building on this previous study we sought to determine the effect of biofilm growth (i) on plasmid persistence in the absence of antibiotics, and (ii) on the evolutionary trajectories of this plasmid in the presence and absence of antibiotics. Our findings can be summarized as follows. First, in the absence of selection, biofilm populations maintained the MDR plasmid longer than planktonic populations. Second, the evolution of plasmid persistence in the absence of antibiotic selection was protracted in biofilms. Third, large deletions of the plasmid conjugative transfer regions were a common evolutionary outcome in planktonic populations but less so in biofilms, indicating that growth in biofilms conserved the plasmid’s ability to transfer horizontally. Finally, plasmid adaptation to one bacterial species can either promote or decrease its persistence in other bacteria, including other opportunistic pathogens.

## RESULTS

### Experimental set-up of biofilm and planktonic populations

To investigate the effects of biofilm growth and the presence of a plasmid-selective antibiotic on the persistence and evolution of MDR plasmid pB10 in *A. baumannii*, we set up an experiment with a full factorial design with two growth environments and two treatments. As shown in Fig. 1, the growth environment was either a biofilm flow cell or serial batch culture; the treatments were the presence (Tet+) or absence (Tet-) of tetracycline in the growth medium. In order to initiate all four environment/treatment combinations with equivalent populations, replicate biofilm populations were first established in flow cells for a period of four days (*t_-4_ – t_0_*). To avoid plasmid loss during this initial phase, the medium was supplemented with Tet for four days, regardless of the subsequent treatment (Tet+ or Tet-). After this biofilm establishment phase at *t*_0_, triplicate flow cells were harvested and used to inoculate triplicate serial batch cultures per treatment. From here on we refer to these serial batch cultures as planktonic populations. The remainder of the growth and sampling scheme is depicted in Fig. 1 (see Materials and Methods for more details).

**Figure 1:**
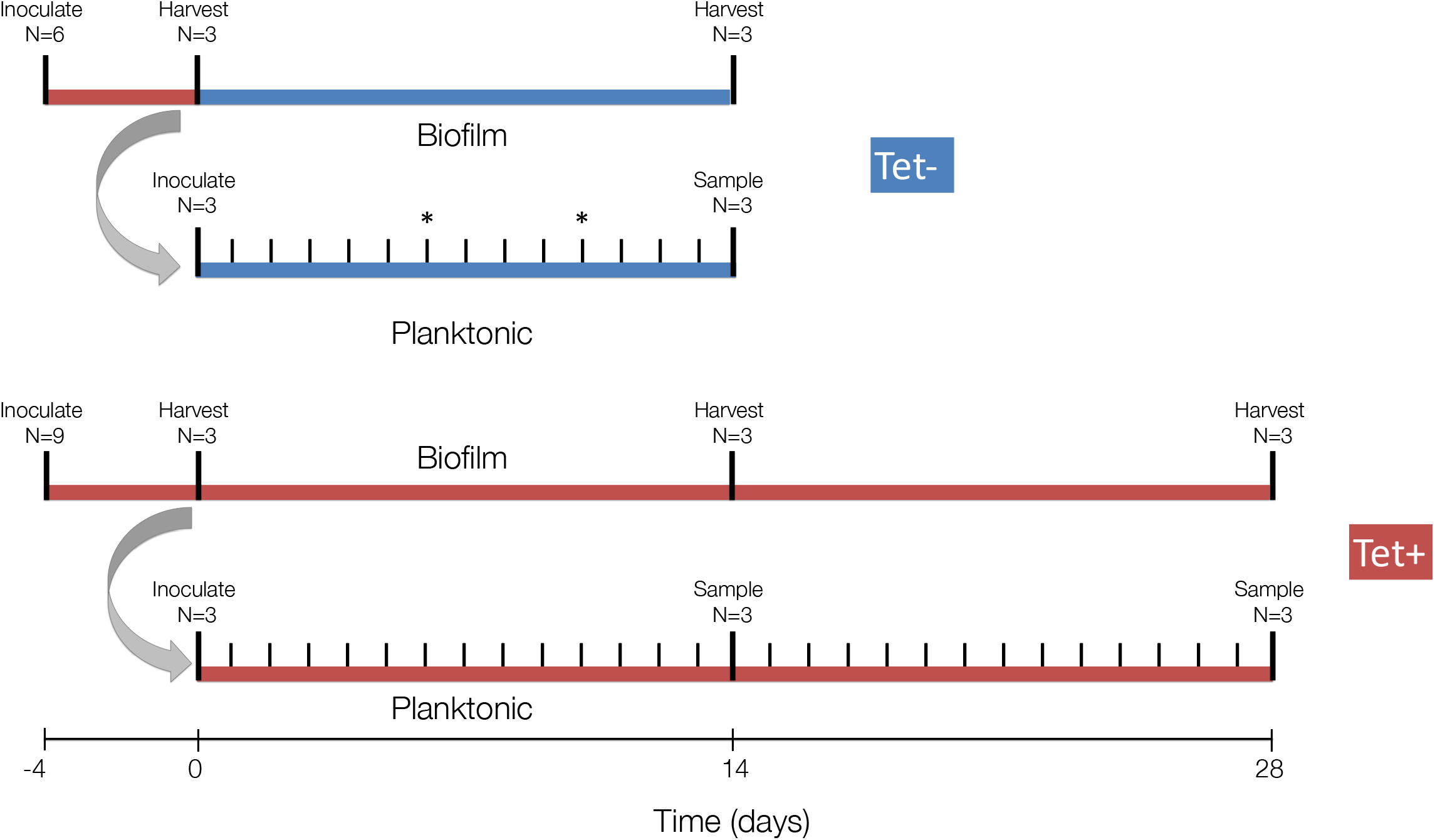
Timeline of the evolution experiments. Red represents time periods when media contained tetracycline (Tet+ treatment); blue represents the absence of antibiotics (Tet-treatment). Large ticks represent inoculation and harvest/sampling events. Four days prior to initiation of the evolution experiments (−4) the biofilm flow cells were inoculated with the ancestral strain. Four days later, at day 0 (*t*_0_), the first set of randomly selected flow cells (n=3) were harvested. A subsample of the cell suspensions from each of these replicates was used to inoculate each of the three planktonic populations (grey curved arrow). Small tick marks indicate the daily serial passage of planktonic populations. Asterisks on days 6 and 10 in the timeline of the Tet-planktonic populations indicate the last day at which plasmid-containing cells were detected in two of the three populations due to rapid plasmid loss; these time points were the final sampling points for these two populations (*t_6_* and *t_10_*). The Tet-experiment was terminated after 14 days and the Tet+ experiment after 28. For more details on the actual protocol, see Materials and Methods and Ridenhour *et al*. ^43^.

### Biofilms retain the plasmid in the absence of selection

To assess the ability of bacterial biofilms to retain an MDR plasmid in the absence of antibiotics, we compared the proportion of plasmid-containing cells present in biofilm and planktonic populations at *t*_0_ and after 14 days of growth in Tet-free media (*t*_14_) (Table 1). This was done by plating samples of these populations on selective and non-selective media. The plasmid was lost more slowly from biofilms than from planktonic populations. After 14 days 12.3 (± 3.5)% of the viable cells from the biofilms still carried the plasmid. In contrast, plasmid-bearing cells could no longer be detected in two of the three planktonic populations (i.e., their frequency in these populations was <10^-8^), and were present in only 0.016% of the third. This observation demonstrates the importance of biofilms in the persistence of plasmid-mediated antibiotic resistance.

**Table 1:** Cell counts as well as absolute counts and fractions of plasmid-containing (P+) cells for each replicate biofilm or planktonic population of the Tet-treatment.

|  | **Total cells**<br>**(per mL)<sup>1</sup>** | **Total viable cells**<br>**(per mL) <sup>2</sup>** | **P+ cells**<br>**(per mL)** | **Fraction P+**<br>**cells <sup>3</sup>** |
| --- | --- | --- | --- | --- |
| ***t0*** |  |  |  |  |
| Rep 1 | 9.6 x 10<sup>9</sup> | 6.1 x 10<sup>6</sup> | 5.9 x 10<sup>6</sup> | 0.98 |
| Rep 2 | 3.9 x 10<sup>9</sup> | 4.7 x 10<sup>6</sup> | 3.8 x 10<sup>6</sup> | 0.82 |
| Rep 3 | 3.5 x 10<sup>9</sup> | 3.4 x 10<sup>6</sup> | 3.4 x 10<sup>6</sup> | 1.00 |
| *t0 Avg* | *5.6 x 10<sup>9</sup>* | *4.7 x 10<sup>6</sup>* | *4.4 x 10<sup>6</sup>* | *0.93* |
| ***t14 Biofilm*** |  |  |  |  |
| Rep 1 | 1.3 x 10<sup>11</sup> | 2.9 x 10<sup>6</sup> | 5.5 x 10<sup>5</sup> | 0.19 |
| Rep 2 | 1.3 x 10<sup>10</sup> | 3.1 x 10<sup>6</sup> | 3.6 x 10<sup>5</sup> | 0.11 |
| Rep 3 | 1.1 x 10<sup>10</sup> | 1.8 x 10<sup>6</sup> | 1.3 x 10<sup>5</sup> | 0.07 |
| *Biofilm Avg* | *5.1 x 10<sup>10</sup>* | *2.6 x 10<sup>6</sup>* | *3.5 x 10<sup>5</sup>* | *0.12* |
| ***t14 Planktonic*** |  |  |  |  |
| Rep 1 | n/a | 2.0 x 10<sup>8</sup> | 3.2 x 10<sup>4</sup> | 0.00016 |
| Rep 2 | n/a | 1.6 x 10<sup>8</sup> | <1 | 0 |
| Rep 3 | n/a | 2.0 x 10<sup>8</sup> | <1 | 0 |
| *Planktonic Avg* | *n/a* | *1.9 x 10<sup>8</sup>* | *1.1 x 10<sup>4</sup>* | *0.00005* |

<sup>1</sup> Total cell counts were obtained by microscopy; <sup>2</sup> total viable cell counts correspond to colony forming units (cfu); <sup>3</sup> P+ fractions were calculated as the ratio of Tet<sup>R</sup> over total cfu counts, obtained by plating on selective and non-selective media. Note that total cell counts estimated by microscopy were only determined for biofilms. n/a: count not performed.

#### Biofilms protracted the evolution of plasmid persistence in the absence of selection

We previously showed that after 28 days of evolution in the presence of Tet, bacteria isolated from biofilms showed a higher diversity in plasmid persistence dynamics than their counterparts evolved in planktonic populations, and on average, a smaller increase in plasmid persistence ^43^. In the current study we compared the evolution of plasmid persistence in clones evolved over 14 days in the presence and absence of Tet (Tet+ and Tet-populations). Because the plasmid was rapidly lost in Tet-planktonic populations, we ended the entire Tet-experiment on day 14. In one of these three Tet-populations, plasmid-containing clones could no longer be detected after day 6, and in another after day 10. Therefore the plasmid-containing clones from Tet-planktonic populations were isolated from samples archived on *t_6_*, *t_10_*, and *t_14_*. In contrast, all the clones from Tet-biofilm populations were isolated on *t_14_*. For each of the triplicate growth environment/treatment combinations plasmid persistence was tested for six clones. First, we verified that clones isolated at *t*_0_ showed the same plasmid persistence profile as the ancestor (Fig. 2A). Based on linear regression there was no significant change in the rate of plasmid loss, meaning the plasmid persistence in the populations used to start the experiment did not change during the 4-day biofilm establishment phase (t-test p-values for *t*_0_, Tet+ was 0.7037 and for *t*_0_, Tet- was 0.5195). Therefore, the *t*_0_ clones were used for the comparisons of plasmid persistence described below.

**Figure 2:**
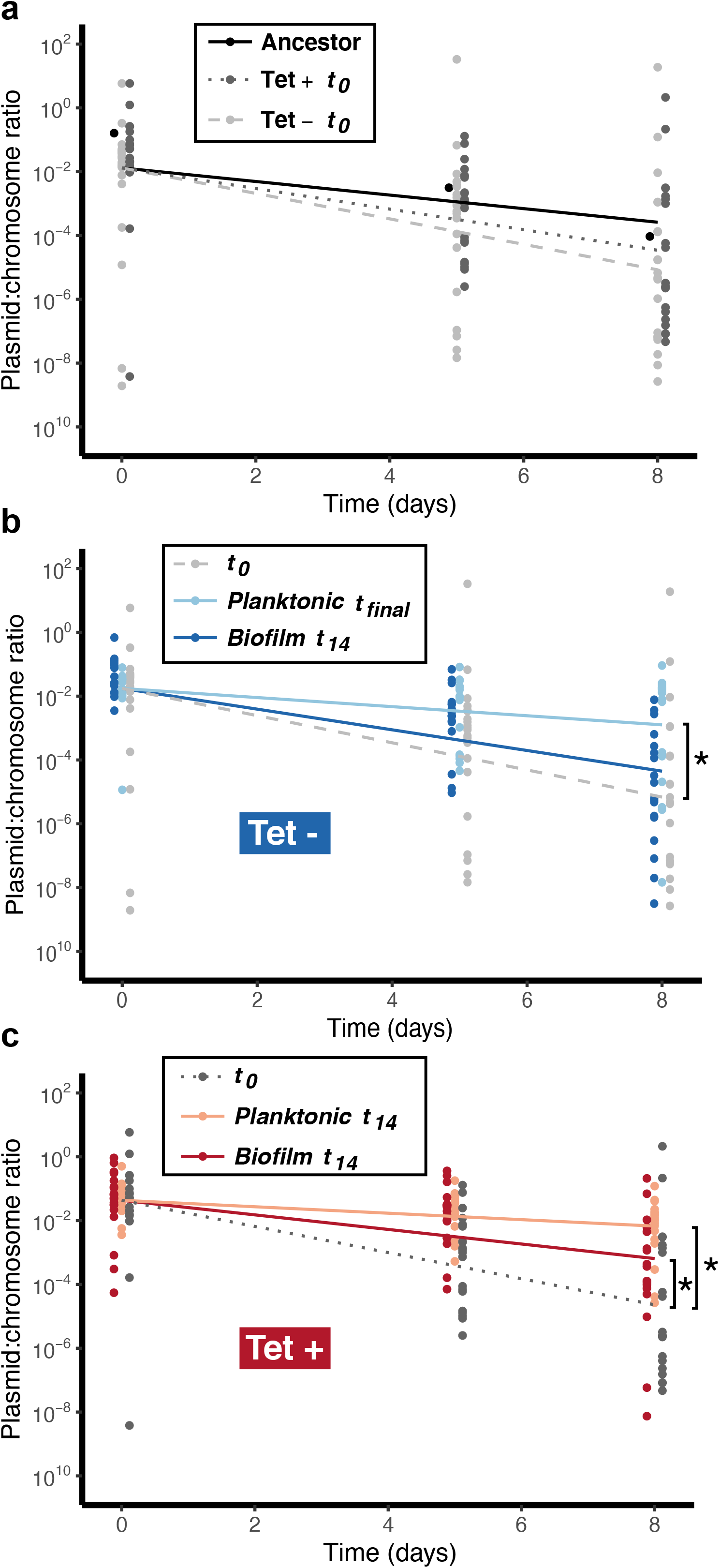
Plasmid persistence shown as the estimated ratio of *trfA*/16S rRNA genes over a period of eight days in liquid serial batch cultures in the absence of selection; (A) the ancestor used to inoculate the flow cells at *t_-4_* compared to clones isolated from all the biofilms harvested after four days of growth with tetracycline, *t_0_*, showing similar plasmid persistence dynamics (Tet*+* and Tet-: populations treated with and without tetracycline after *t_0_*); (B) clones isolated from the Tet-populations, showing a drastic increase in plasmid persistence after evolution in planktonic but not in biofilm populations (note that two of the three planktonic populations no longer contained detectable plasmids after day 6 and 10, respectively, requiring analysis on these days instead of *t_14_* – the final sampling times are therefore referred to as *t_final_*); (C) clones isolated at *t_0_* and *t_14_* of the Tet+ evolution experiment, showing an increase in plasmid persistence over time for all populations. Lines are the output of the log-linear model for each group. * denotes a significant difference (t-test p-value <0.05) between the groups pointed out by the brackets: planktonic, *t_final_* versus *t_0_* and biofilm *t_14_* versus *t_0_*. The plasmid persistence data from *t_0_* in the Tet+ experiment were reported by us previously ^43^.

After 6–14 days of evolution in the absence of antibiotics, plasmid pB10 showed improved persistence in *A. baumannii* clones from planktonic populations but not in clones from biofilms (Fig. 2B; t-test p-values were 0.0007 and 0.2107, respectively). These results are in contrast to the clones evolved in the same conditions but in the presence of Tet, where plasmid persistence improved in clones isolated from both biofilm and planktonic populations after 14 days (Fig. 2C; t-test p-values were 0.0053 and <0.0001, respectively). The latter result is consistent with our previously published findings after 28 days of biofilm and planktonic growth ^43^. Thus on average, plasmid persistence always improved in planktonic populations regardless of antibiotic treatment, but only improved in biofilms grown with Tet.

### Plasmid transfer genes were mostly retained in biofilms but completely lost in planktonic populations

To determine the genotypic changes that occurred during the experiment, we sequenced all 18 clones per environment/treatment at each time point (6 per triplicate population, Fig. 1), for a total of 144 strains. Here we focus on genetic changes in the plasmid pB10. We identified three types of mutations, as shown by a visual summary in Fig. 3 and a list in Supplementary Table 1. By far the most common genetic changes were large deletions in the plasmid regions encoding conjugative transfer (*tra*), mating pair formation (*trb*), and an intervening class 1 integron containing sulfonamide and amoxicillin resistance genes. These large deletions also often included a few genes thought to be involved in plasmid maintenance and central control, adjacent to the *tra* region. Conjugation assays with a representative clone showed that this gene loss resulted in the inability of these plasmids to transfer by conjugation. When the donor strain contained a deletion mutant of plasmid pB10 we could not detect any transconjugants (Supplementary Table 2). In contrast, transconjugants were detected for the ancestor and a randomly chosen evolved clone with intact plasmid, and the average counts (T) were significantly different from 0 (t-test p-values were 0.03662 and 0.01467, respectively).

**Figure 3:**
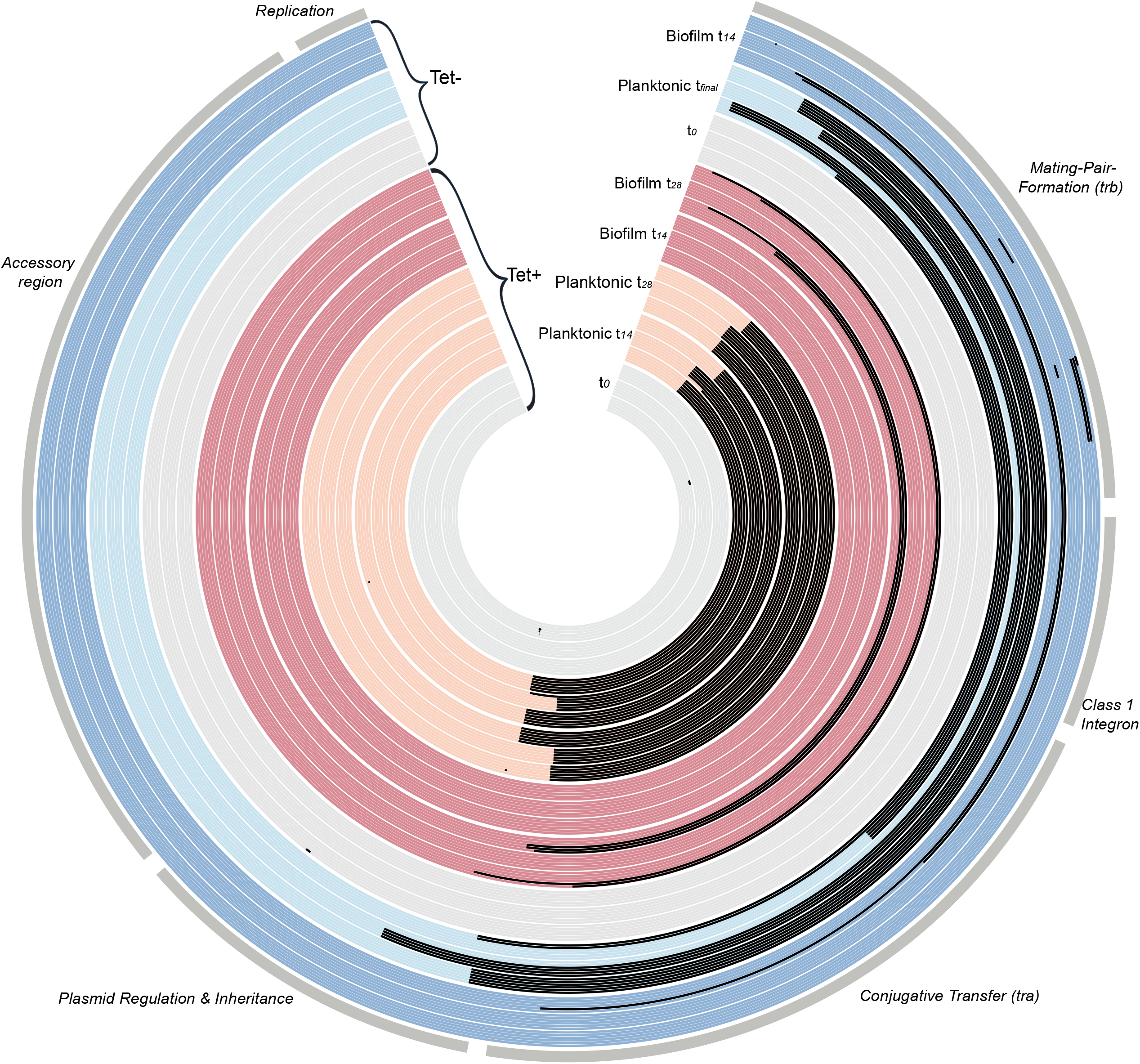
Maps of evolved plasmids with major segment functions identified, showing all plasmid mutations observed in each sequenced clone evolved under different treatments (Tet+ and Tet-) in different environments (biofilm and planktonic populations). Evolved clones are grouped by treatment, environment, and time point, as shown by different colors and shades and identified in the open wedge of the circle. The plasmid genome of each clone is shown as a single band – at each time point, 18 clones were analyzed per treatment and environment (six clones from each triplicate population). Tet+, Tet-: evolved with and without Tet. Black areas in the plasmid maps represent deletions (note that small indels are shown as small spots) and the dots represent SNPs; see Supplementary Table 1 for details. Note that the tetracycline resistance operon is located in the ‘accessory region’.

We also identified nine deletions smaller than 1,000 bp located in maintenance/control genes *kfrA*, *klcB*, or one of the *trb* genes. Finally, there were four single nucleotide polymorphisms (SNPs): two in *kfrA*, one in a *trb* gene, and one in an intergenic region near the *tetR* gene. In summary, large deletions that remove virtually all the genes required for horizontal transfer were by far the most common genetic changes.

The size of the large deleted plasmid regions varied drastically between clones, ranging from 21,755 bp to 34,594 bp. The number of genes deleted per plasmids ranged from 23 to 35, but a common set of 19 genes (from *trbF* through *traC*) were deleted or truncated in every deletion variant (Fig. 3, Supplementary Table 1). As plasmid pB10 has 65 complete coding sequences (CDS) in addition to a few truncated transposases ^53^, all large deletions represented more than a third of the gene content of the plasmid. Whereas the boundaries of these deletions were all different, most contained short flanking direct repeats (typically 8-12 bp, with one being only 6 bp, Supplementary Table 1). One of these repeats was always eliminated during deletion of the intervening region. Deletions between these direct repeats are indicative of recombination events ^54^.

There were striking differences in the kinds of plasmid genotypes observed in the biofilm versus planktonic populations (Fig. 3, Supplementary Table 1). Most of the clones from planktonic populations showed the large deletions described above. Only 2 of the 54 clones had retained the ancestral pB10 sequence, both from the Tet-population sampled at *t*_6_. In five of the six Tet+ planktonic populations sampled at either 14 or 28 days, all six clones contained plasmids with identical large deletions. Moreover, two of the three planktonic populations showed the same large deletion at both time points. These results indicate strong selective sweeps of large deletion mutants in planktonic populations under antibiotic selection for the plasmid. Even in the absence of Tet, all six clones from the population that still retained plasmids by day 14 showed an identical large deletion. In stark contrast, all clones from the three Tet+ biofilms at *t*_14_ still contained the ancestral plasmid genotype. Even two weeks later, one of the three Tet+ biofilm populations still showed intact plasmids in 100% of the clones, and in the other two populations around half the clones still had intact plasmids. In the Tet-biofilms the deletions were more frequent at *t*_14_ but still only found in half or less than half of the clones per population (Supplementary Table 1 and Fig. 3). Moreover, of the 54 evolved biofilm clones only two pairs showed an identical deletion. These results clearly show that there were no sweeps of large deletion mutants in the biofilms. In conclusion, biofilms still contained a large proportion of transmissible plasmids after 14 or 28 days, whereas transmissible plasmids were no longer detected at these time points in planktonic populations.

### Loss of plasmid transfer genes improved plasmid persistence

To determine the effect of the large plasmid sequence deletions, we compared the persistence of truncated and full-length plasmids in two ways. First, we compared the persistence of these plasmids in their co-evolved hosts. On average, clones with truncated plasmids showed a significantly higher plasmid persistence than clones with full-length plasmids (Fig. 4A, t-test p-value was < 0.0001). However, a few biofilm clones with full-length plasmids identical in sequence to ancestral pB10, also demonstrated high plasmid persistence (Fig. 4A), suggesting this change was caused by chromosomal mutations. Next, to ensure that the observed improvement in persistence of the truncated plasmids could entirely be explained by the plasmid deletions and not chromosomal changes, we transformed the ancestral host with one full-length pB10 and several truncated plasmids from *t*_28_ of the Tet^+^ treatments. Plasmid persistence was higher for the truncated plasmids than for the full-length plasmid (t-test p-value was < 0.0001) or ancestral pB10 (Fig. 4B; t-test p-value was 0.0151). Thus, the large deletions in plasmid pB10 were a major driver of improved plasmid persistence in *A. baumannii*.

**Figure 4:**
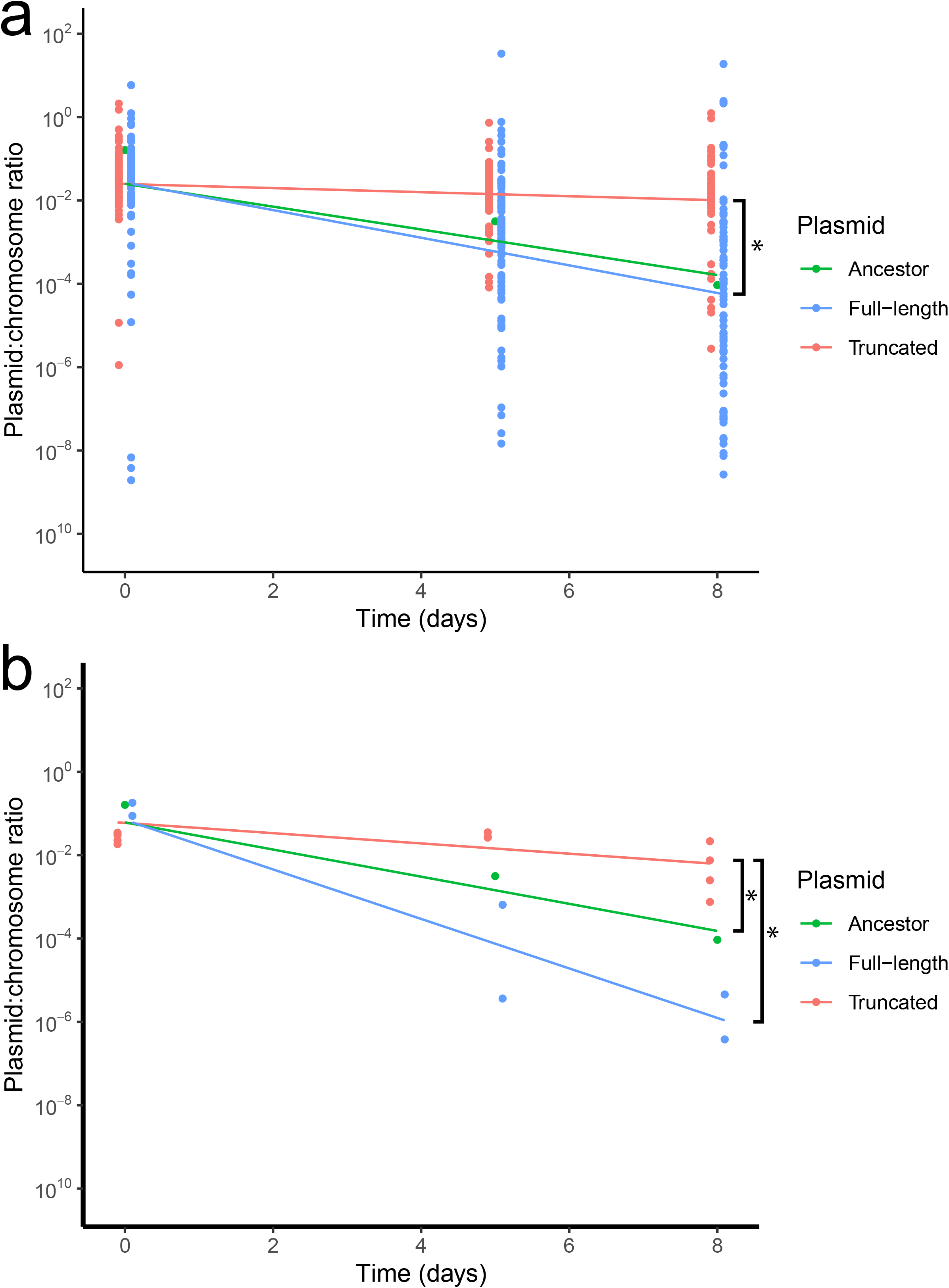
Persistence of plasmids evolved in Tet+ biofilms and planktonic populations tested in (A) their co-evolved hosts (92 full-length and 49 truncated plasmids), and (B) in the ancestral host (2 full-length and 4 truncated plasmids). In both panels the ancestral plasmid was tested in the ancestral host. Plasmid persistence is shown as the estimated ratio of *trfA*/16S rRNA genes over time. Lines are the results of the log-linear model for each group. * denotes a significant difference (t-test p-value was <0.05) between the groups indicated by the brackets: truncated plasmids versus ancestral plasmid and versus full-length plasmids.

#### Persistence of evolved pB10 in other naive hosts

To understand the broader effects of the large deletions in pB10 on plasmid persistence, we tested one evolved truncated plasmid from a *t*_28_ clone in three bacterial strains: *Pseudomonas moraviensis* R28, *Pseudomonas sp. nov*. H2, and *Stenotrophomonas maltophilia* P21 (see footnote 3 in Supplementary Table 1). All three strains were previously shown to rapidly lose pB10 ^55^. The evolved plasmid was even less persistent than ancestral pB10 in the two *Pseudomonas* hosts, but it was more persistent in *S. maltophilia* P21 (Fig. 5). Our data demonstrate that even though deletions that eliminate conjugative transfer of the plasmid may be beneficial in some species, that benefit may not extend to all.

**Figure 5:**
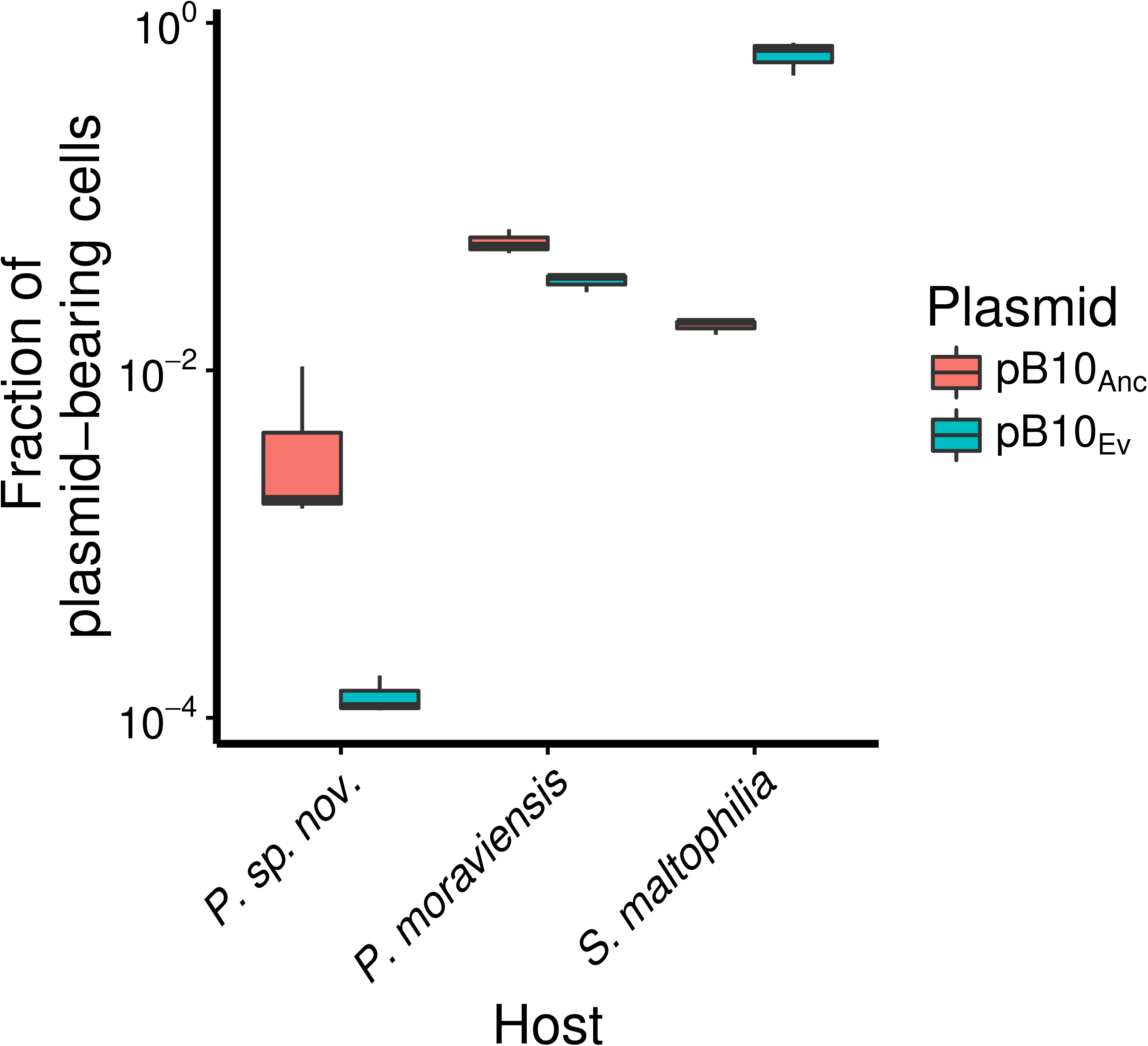
Plasmid persistence of ancestral plasmid pB10_Anc_ and truncated plasmid pB10_ev_ in three additional hosts, as determined by plate counting after 8 days, showing that the plasmid deletion was only beneficial to persistence in *Stenotrophomonas maltophilia*. (P= *Pseudomonas*). Boxes represent the median and the first and third quartiles, and the whiskers the lower and upper 1.5 interquartile range.

### Some chromosomal mutations are also responsible for improved plasmid persistence

While the mutations found in the plasmids explained most of the increased persistence in the isolated clones, we identified a few clones where chromosomal mutations must be responsible for the improved persistence (Figure 4). In the genome sequences of the 144 clones, we identified many chromosomal mutations when compared to the ancestral sequence (Supplementary Table 3). However, none of these mutations were in genes previously shown to be involved in plasmid persistence. Due to the high number of unique mutations, mutation reconstruction in the ancestor would be required to test their effects, which is outside the scope of this study.

## DISCUSSION

In this study we showed that growth in biofilms can promote the retention of MDR plasmids, while preserving the horizontal mode of plasmid transmission better than planktonic populations. These findings have several important implications. First, they demonstrate the positive effect of biofilm growth on MDR plasmid persistence in the absence of antibiotics. This is significant given most bacteria in clinical and environmental habitats grow in biofilms ^25^. Our findings are consistent with previous studies that showed that the deeper layers of biofilms can act as refugia for plasmids even in the absence of selection for the plasmid ^56,57^. A plausible explanation is that bacteria in biofilm layers farthest away from the bulk medium grow slowly or not at all. This allows plasmids to be better retained in these subpopulations as plasmid loss requires cells to divide. Our results thus suggest that reduced use of antibiotics may help lower the relative abundance of resistance genes in biofilms but not necessarily eliminate them from a population. Second, evolution in biofilms conserved the horizontal mode of transmission of a MDR plasmid, while growth in well-mixed systems resulted in sweeps of much smaller, non-self-transmissible plasmids that persist better through vertical transmission. Finally, this study cautions against indiscriminately extrapolating results from experimental evolution in planktonic cultures to more complex environments such as bacterial biofilms.

Perhaps the most important finding of our study was the striking difference in evolutionary trajectories between the biofilm and planktonic populations. Independent of the presence of selection for the plasmid, virtually every tetracycline resistant *A. baumannii* clone analyzed from planktonic populations had lost a very large segment of plasmid pB10 that contained horizontal transfer and a few antibiotic resistance genes. The loss of these genes resulted in the inability of the plasmid to transfer by conjugation (Supplementary Table 2). While such gene loss may not be a general adaptive strategy for most conjugative plasmids, the findings are consistent with other studies that observed loss of plasmid transmissibility due to deletion or mutations of transfer genes during evolution in planktonic populations ^21,44,58,59^. In contrast, the majority of plasmids in evolved biofilm clones remained intact and thus self-transmissible. In only one of the nine evolved biofilm populations did plasmid genotypes with deletions in transfer genes outnumber the ancestral genotype (a Tet+ biofilm at *t*_28_). Biofilms can thus protract the emergence of non-transmissible plasmid mutants.

There are a few possible explanations for these contrasting results between biofilms and planktonic populations. The most parsimonious one is that the spatial structure of biofilms protracted selective sweeps, which is in line with previous studies ^29–33,44^. Although we did not compare plasmid fitness cost, the higher persistence of the much smaller evolved plasmids can almost certainly be explained by a reduced fitness cost on their host. First, loss of the two transfer regions would be expected to reduce plasmid cost as horizontal transfer (conjugation) is known to be costly ^21,58,59^. Moreover, in the class 1 integron region that was always deleted together with the transfer regions, the integrase gene could have added to the cost of the plasmid ^60,61^. Since this gene is not repressed by LexA in *Acinetobacter* spp., a highly active integrase expressed from pB10 could have had a toxic effect on *Acinetobacter* spp. ^62^. Regardless of which genes conferred the highest cost, losing these large plasmid segments allowed the hosts of truncated plasmids to rapidly sweep through the planktonic Tet resistant populations. In contrast, due to the absence of global competition in biofilms, clones with wild-type plasmids were not as readily outcompeted by clones with deletion variants. An argument could be made that the bacteria in biofilms may have undergone fewer doublings than those in planktonic populations ^63,64^, thus slowing evolution. Unfortunately, while the number of doublings in the planktonic populations can readily be estimated, this is not the case for biofilm populations where the growth rate varies according to depth and the availability of nutrients. Even if the number of generations differed between the two environments, what matters is that within a certain time frame, such as a course of antibiotics, some biofilm forming pathogens may lose the ability to transfer their MDR plasmids less rapidly than expected based on results from well-mixed systems.

An additional hypothesis that can explain the increased retention of plasmid transfer genes in bacteria evolving in biofilms is that a tradeoff exists between horizontal and vertical transmission ^58^. It is known that in biofilms close cell-to-cell contact and stabilized mating pair formation can improve the efficiency of conjugative transfer to neighboring cells relative to planktonic populations ^65–68^. This makes horizontal transfer a more important mechanism of plasmid ‘survival’ in biofilms ^69^. It would then follow that selection on plasmid transferability may be stronger in biofilms than in axenic planktonic populations, where it would present a cost rather than a benefit. This is especially true for IncP-1 plasmids like pB10, which do not transfer efficiently in shaken liquid cultures due to their short rigid sex pili ^70,71^. In such a well-mixed environment, smaller non-self-transmissible plasmids with a lower fitness cost could thus outcompete transmissible ones and propagate through vertical transmission only.

Another plausible explanation for higher persistence of self-transmissible plasmids in biofilms is the positive selection for retention of the sex pilus genes in this environment, which are part of the transfer gene region *trb* that was deleted in the planktonic clones. Previous studies have shown that conjugative plasmids can stabilize biofilms during the early stages of biofilm formation due to the formation of sex pili ^65,72^. If pilus expression was favorable for biofilm formation in our flow cells, loss of the pilus could have been selected against in this environment.

Overall, our results suggest that in planktonic populations, the cost of maintaining the transfer regions and integron is apparently higher than the benefit of retaining the plasmid through horizontal transfer. The stark difference in plasmid evolutionary trajectories between planktonic and biofilm populations and the higher diversity in plasmid persistence phenotypes reported by us previously ^43,44^ suggests that it is critical to study bacteria-plasmid coevolution in biofilms to fully understand plasmid population biology.

The deletions of the large plasmid segments in *A. baumannii* may result from homology-facilitated illegitimate recombination due to short direct repeats that flanked the boundaries^54^. Illegitimate recombination due to short repeats is well understood in *E. coli*, and has also been reported in *Acinetobacter baylyi* ^73^. The same plasmid pB10 was previously shown by us to lose its tetracycline resistance operon in *E. coli* through recombination between much larger direct repeats ^74^. Interestingly Porse *et al*. (2016) ^21^ identified a similar pattern of large deletions of a plasmid’s conjugative transfer region in an IncN plasmid from *Klebsiella pneumoniae* that was evolved in strains of *Escherichia coli*. In that study the observed deletions were very similar to one another and corresponded to the location of IS26, whereas the pB10 deletions in our study were not adjacent to any IS element. Given the very short length of the flanking repeats on plasmid pB10 and the very large deleted fragments, it would be interesting in future studies to determine the rate at which these deletions occur in this *A. baumannii* strain.

In our previous studies we never observed loss of the entire plasmid transfer regions of pB10 or other plasmids of the same IncP-1 group ^13,17,18^. Therefore we examined the persistence of one of our pB10 deletion variants in three other bacterial species. The deletion caused a large increase in plasmid persistence in *S. maltophilia*, but not in two environmental *Pseudomonas* isolates. *S. maltophilia* is increasingly found to be an important nosocomial pathogen^75^. Our findings thus show that deletion of plasmid transfer genes could also contribute to the persistence of MDR plasmids in distantly related bacterial pathogens.

The persistence of plasmids that remained in the planktonic populations had always improved within 14 days (140 generations) or less, regardless of the presence or absence of Tet. This finding is consistent with previous work that showed a reduction in plasmid cost or improvement of persistence over time in the absence of plasmid-selective antibiotics^15,19,76^. Thus even when MDR plasmids are in bacteria that are not exposed to antibiotics, they can undergo genetic changes that rapidly improve their persistence. Upon later exposure to antibiotics, these deletion variants would sweep through the population, as they can outcompete ancestral or plasmid-free clones. In contrast, in biofilms the average plasmid persistence only improved significantly in the Tet+ biofilms. The observed discrepancy between biofilm and planktonic populations may be due to the same variety of factors described above.

Our findings suggest that bacterial biofilms play an important role in the maintenance and spread of MDR plasmids in natural bacterial populations in several ways. First, growth in biofilms can increase MDR plasmid persistence in the absence of antibiotics. Second, evolution in biofilms may facilitate the preservation of plasmid transferability. So far this has not been considered as a possible positive effect of biofilm growth on the spread of MDR plasmids. These findings point out that while experimental evolution of bacteria in planktonic populations have addressed important basic evolutionary questions, these growth conditions are far removed from those of naturally occurring bacterial populations. It is thus vital that future research on the ecology and evolution of MDR plasmids includes studies on bacterial biofilms.

## METHODS

### Bacteria, plasmid, and growth media

The ancestor used in our study was derived from *Acinetobacter baumannii* strain ATCC 17978 (Accession #CP000521) and is the same as used in our previous study ^43^. Here we refer to this strain simply as *A. baumannii*. The plasmid used was pB10, a 64.5-kbp broad-host-range IncP-1 plasmid from a German wastewater treatment plant ^53^ that is poorly maintained in naïve *A. baumannii*. It encodes resistance to tetracycline, streptomycin, amoxicillin, sulfonamides, and HgCl_2_. Upon whole-genome sequencing of the ancestral strain after electroporation with pB10 we observed the loss of a large DNA segment (~140-kbp). This region was previously thought to be a chromosomal island ^77^ but more recently shown to be part of a 150-kbp plasmid pAB3 in this strain ^78^. We determined that upon introduction of plasmid pB10 in ATCC 17978 by electroporation, our ancestor lost pAB3 and is thus a derivative of ATCC 17978 without that mobile genetic element. In addition to pB10, the strain retained the two small native plasmids, pAB1 and pAB2. Other strains used in this study were *Pseudomonas* sp. nov. H2 ^79^, *Pseudomonas moraviensis* R28 ^80^, and *Stenotrophomonas maltophilia* P21 ^55^.

For information regarding the construction of the ancestral host, the MBMS medium with and without tetracycline (Tet, 10 mg/l), and the culture conditions, we refer to Ridenhour et al. ^43^.

### Experimental evolution protocol

A timeline of our experimental plan is shown in Fig. 1. The evolution experiment consisted of a full factorial design with two growth environments and two treatments. The growth environment was either a biofilm flow cell or serial batch culture. In one treatment (Tet+) the plasmid-selective antibiotic Tet was added during the entire length of the experiment, whereas in the Tet-treatment Tet was only present during the 4-day biofilm establishment period (*t_-4_ – t_0_*). This antibiotic selection phase was necessary to avoid establishing biofilms that would consist of a mixture of plasmid-containing and plasmid-free cells. In all treatments, time point *t*_0_ represents the end of the ‘biofilm establishment phase’, which occurred four days after the flow of MBMS-Tet through the flow cells was initiated following strain inoculation. As the set-up and growth conditions of the Tet+ treatments have been described in detail by us previously ^43^, we focus here on conditions that differed between treatments. The ancestral strain, grown overnight in MBMS-Tet, was inoculated in multiples of three flow cells, as biofilm sampling was destructive, and triplicate populations were harvested at each time point. This harvesting was done at *t*_0_ and after 14 and 28 days (*t*_14_, *t*_28_) for the Tet+ treatment, and at *t*_0_ and *t*_14_ for the Tet-treatment. The Tet-treatment was ended at *t*_14_ instead of *t*_28_, due to rapid plasmid loss in the parallel Tet-planktonic populations. The harvesting procedures are detailed in Ridenhour *et* al. ^43^.

The Tet+ and Tet-planktonic populations were initiated by inoculating three replicate test tubes containing five ml of MBMS-Tet or MBMS with cell suspensions from the triplicate *t*_0_ biofilms of the Tet+ and Tet-treatments, respectively (Fig. 1). Each of these planktonic populations were then grown in parallel to the remaining biofilm populations. We used this inoculation approach to control for the level of diversity already present in the biofilms at *t*_0_, due to evolution that had taken place during the biofilm establishment phase. Following inoculation, the planktonic populations were serially passaged every 24 (± 1) hours to obtain approximately 10 generations per day, as described previously ^43^. For the Tet+ treatment, the planktonic populations were archived at *t*_14_ and *t*_28_. For the Tet-treatment, the planktonic populations were archived daily because we did not know how long plasmid-containing (Tet-resistant) cells would be maintained in these populations. Appropriate dilutions were plated at *t*_5_, *t*_10_, and *t*_14_ on lysogeny broth agar (LBA) and LBA supplemented with tetracycline (10 mg/l) (LBA_Tet_). When plasmid-containing cells were no longer detected on LBA_Tet_, the populations archived at previous time points were plated until we could identify the last time point at which the population still contained plasmid-bearing cells.

For both biofilm and planktonic populations at each sampling point (see Fig. 1), serial dilutions of cell suspensions were plated on LBA_Tet_ and plates were incubated overnight. To isolate evolved clones for plasmid persistence tests and whole-genome sequencing, six resistant clones per replicate population were purified by restreaking twice and archived. The procedures used for archiving clones and for plasmid extractions were described by us previously ^43^.

To determine the effect of large plasmid deletions on plasmid persistence, we also transferred plasmid DNA extracted from five resistant clones obtained from *t*_28_ of the Tet+ treatment into the ancestral *A. baumannii* host. This was done by electroporation, essentially as described previously, except for washing the cells in cold deionized water^81^.

### Plasmid fate in biofilms versus planktonic populations in the absence of antibiotics

To determine the loss of the plasmid in the Tet-biofilms and planktonic populations between *t*_0_ and *t*_14_, cell suspensions were diluted and plated on LBA and LBA_Tet_. From these colony-forming unit (cfu) counts, we determined the total number of culturable cells and the proportion of plasmid-containing cells. In addition to the number of culturable (=viable) cells, we also determined the total number of cells in the Tet-biofilms using phase-contrast microscopy and a Petroff-Hausser counting chamber.

### Plasmid persistence assays

To determine plasmid persistence, we performed triplicate plasmid persistence assays on six clones per time point per treatment, as described in Ridenhour *et al*. ^43^. Briefly, each clone was grown overnight in MBMS-Tet and used to inoculate three test tubes with MBMS, which were incubated for 24 hours (+/- 1 hour) before passage into new media. This was repeated for a total of 8 days. To estimate the fraction of plasmid-bearing cells on days 5 and 8 by real-time quantitative PCR (qPCR), cell pellets obtained after centrifugation of population samples were frozen at −20°C. DNA extractions and qPCR reactions were performed to determine the copy number of the pB10-encoded replication initiator gene *trfA* relative to the number of 16S rRNA copies. These qPCR-based estimates of the plasmid:chromosome ratio were used as a measure of the fraction of plasmid-bearing cells. This gene ratio is a proxy for the fraction of plasmid-containing cells but not an exact representation, as there are five copies of the 16 rRNA gene in *A. baumannii* strain ATCC 17978 and on average three copies of the plasmid pB10 (Supplementary Figure 1). In addition, this ratio may be slightly affected by slight variations in copy number (i.e., less than two-fold) of the pB10 and partial chromosomes replicating in fast-growing cells. However, these variations were small, as Supplementary Fig. 1 shows very limited variation of the plasmid:chromosome sequence read coverage among all sequenced clones. Given this, the significant decline of plasmid:chromosome ratios in some populations during the plasmid persistence assays was not due to changes in plasmid copy number per cell, but due to plasmid loss in these populations. As described by us previously ^43^, we used the log-linear model to estimate the rate of plasmid loss over time.

### Plasmid persistence in other hosts

We extracted ancestral pB10 and one evolved truncated pB10 variant from a clone obtained from a *t*_28_ biofilm (Supplementary Table 1) and used electroporation to insert each into three additional bacterial strains. Plasmid persistence assays for these clones were performed as described above, but for a total of seven days. They were sampled at days 0, 4, 7, and plasmid presence was determined using serial dilutions and plate counting on LBA and LBA_Tet_ instead of qPCR.

#### Plasmid transfer assays

The ability to transfer their plasmid by conjugation was compared for evolved and ancestral strains as described by us previously, with *E. coli* K12Nal as the recipient ^17^. Briefly, 1 ml of donor and recipient cultures – grown overnight at 37 °C were harvested by centrifugation and resuspended in 100 μl LB broth; 50 μ l of each cell type was then mixed together and spotted onto an LB agar plate for the actual conjugation and spotted separately as negative controls. Mating plates were incubated at 37 °C. After overnight incubation, the entire cell mass of each control and conjugation mixture was harvested and suspended in 500 μl saline, and a dilution series were spread onto the appropriate donor-, recipient- and transconjugant-selective LB agar media.

### Sequencing, and sequence analysis

For DNA sequencing, total genomic DNA from each clone was extracted from 2mL of an overnight culture using the GenElute Bacterial Genomic DNA kit (Sigma-Aldrich) and the quality and integrity of the DNA was assessed on a 1% agarose gel and the concentrations were determined fluorometrically using Quant-iT™ PicoGreen^®^ dsDNA Reagent (ThermoFisher Scientific, Waltham, MA, USA) with the TBS-380 Mini-Fluorometer (Turner BioSystems) (Molecular Devices, Sunnyvale, CA, USA). Samples were submitted to the University of Idaho IBEST Genomic Resources Core facility for library preparation and whole-genome sequencing using Illumina MiSeq (Illumina, San Diego, CA, USA) and associated chemistry.

Following sequencing, the sequence data were screened to remove low-quality reads, sequencing adapters, and duplicate read pairs using the software package htstream (https://s4hts.github.io/HTStream/). To identify mutations in the genome, cleaned reads were mapped against the reference sequence for pB10 (AJ564903.1) and *A. baumannii* ATCC 17978 (GCA_000015425.1) using *breseq* v0.35.7 ^82^. Prior mapping discrepancy between the reference sequence of *A. baumannii* ATCC 17978 and our ancestral seed stock were corrected using the gdtools APPLY command from *breseq*. The command was applied iteratively until no more mutations were detected between the ancestor reads and the reference sequence. Mutations between strains were compared using the gdtools COMPARE from *breseq* and the presence of undetected deletions was manually screened by searching for Missing Coverage Evidence and the presence of corresponding New-Junction Evidence, indicating reads spanning the area where no reads were mapped. The plasmid copy numbers shown in Supplementary Figure 1 were estimated using the predicted mean read-depth coverage given by *breseq* for the plasmids and the chromosome (note that the predicted values are corrected for the effects of deleted regions).

### Statistical analysis

Statistical analysis of the qPCR data was done using a linear mixed-effect model approach in R with the nlme package as detailed in Ridenhour *et al*. ^43^. Briefly, the log plasmid:chromosome ratio of each group was predicted using the origin of the clones (i.e., *t*_0_, biofilm, or planktonic population) as fixed effect and the clone replicate measure as random effect. Each parameter was nested within the day of the experiment (Days 0, 5, 8) as a continuous variable.

## DATA AVAILABILITY

All sequencing data pertaining to this project have been made available at the National Center for Biotechnology Information (SRA accession number PRJNA869051). All plasmid persistence data are available at https://github.com/tstal/Metzger-et-al-2022.

## CODE AVAILABILITY

The code used analyze the plasmid persistence data is available at https://github.com/tstal/Metzger-et-al-2022.

## ACKNOWLEDGEMENTS

This project was funded by grant DOD-DM110149. G.A.M. was also in part supported by the Bioinformatics and Computational Biology Program at the University of Idaho in partnership with the Institute for Bioinformatics and Evolutionary Studies (now Institute for Interdisciplinary Data Sciences, IIDS). T.S. and E.M.T. were also partially supported by the National Institute of Allergy and Infectious Diseases at the National Institute of Health (grant number R01 AI084918). The genome sequencing was carried out by the IIDS Genomic and Bioinformatic Resources Core, and made possible thanks to NIH National Institute of General Medical Sciences Award No. P30 GM103324. We would like to thank Wesley Loftie-Eaton, and Silvia Smith for helpful discussion on the design of this project, and Rachana Regmi, Andrew Avery, Sean West and Taylor Wilkinson for their roles in data collection.

## COMPETING INTERESTS

The authors declare no competing interests.

## AUTHOR CONTRIBUTIONS

E.M.T. and L.J. F., conceived the project; G.A.M., E.M.T., L.J. F., B.J.R., and M.L.S., designed the study; G.A.M., M.F., K.G., and J.M. performed experiments and collected the data; G.A.M. and T.S. performed the genomic analysis; G.A.M. and B.J.R. performed the statistical analyses; G.A.M., T.S. and E.M.T. wrote the manuscript; All authors helped revising the manuscript.

## Notes

https://github.com/tstal/Metzger-et-al-2022

### Competing Interest Statement

The authors have declared no competing interest.

### Summary of Updates

The paper was revised in response to three reviewers' comments. It will be accepted by the journal in the near future.

