## Supplemental figure and tables for "Biofilms preserve the transmissibility of a multi-drug resistance plasmid"

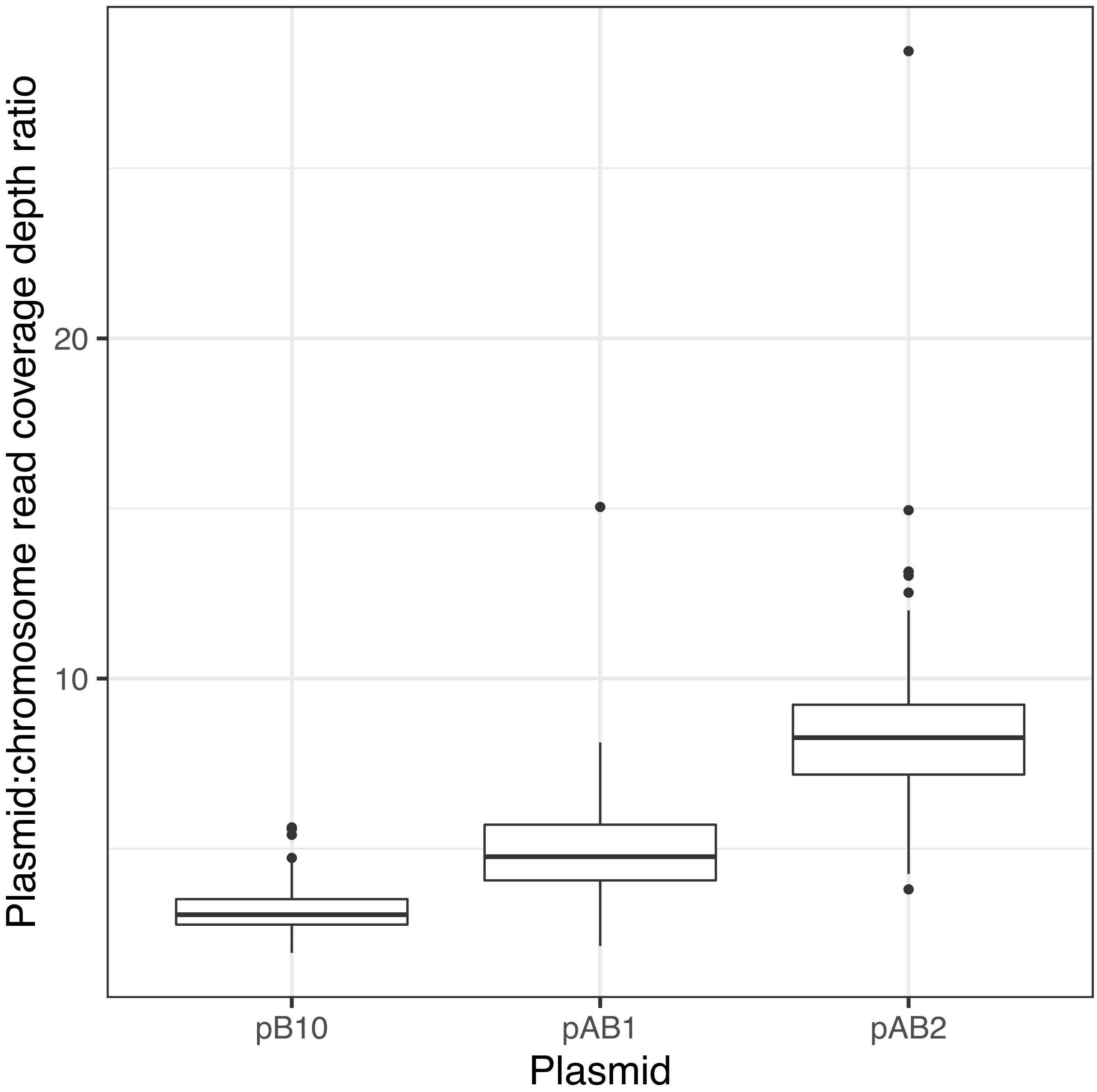


Supplementary Figure 1: Plasmid copy number of the ancestral and all evolved clones based on plasmid-chromosome read coverage for all whole genome sequences.

**Supplementary Table 1: Summary of evolved plasmid genotypes with their affected genes and functions, found in all 144 sequenced clones**

| Antibiotics | Environment | Time Point | Gene(s) | Mutations | Gene Function | Population Replicate | Direct Repeats that Flank Deletions (5'-3'), if present |
|---|---|---|---|---|---|---|---|
| **Tet+** | Starting Point | $t_0$ | *kfrA* | Δ17 bp | KfrA protein in maintenance/control region | C | |
| | | | " " | Δ141 bp | " " | C | |
| | | | " " | 2 SNPs | " " | C | |
| | | | *trbL* | Δ405 bp | putative DNA topoisomerase in transfer region | C | |
| | Biofilm | $t_{14}$ | No plasmid mutations at all | | | | |
| | | $t_{28}$ | *trbD–kfrA* | Δ33,164 bp | 34 genes in transfer and maintenance/control regions | A | GGCCTG |
| | | | *trbB–traL* | Δ31,764 bp | 32 genes in transfer region | A | |
| | | | *trbE–kfrC* | Δ30,220 bp (x2) | 31 genes in transfer and maintenance/control regions | C | GGCGGCCTCC |
| | | | *trbB–kfrC* | Δ32,730 bp | 34 genes in transfer and maintenance/control regions | C | CGCGGACGGC |
| | | | *trbJ* | Δ9 bp | conjugal transfer protein TrbJ | C | |
| | Planktonic | $t_{14}$ | *trbF–kfrA* | Δ29,446 bp (x7) | 32 genes in transfer and maintenance/control regions | A, B | GCGCGCGC |
| | | | *xf2080 / tetR* [2] | SNP | hypothetical protein / tetracycline resistance repressor | A | |
| | | | *trbE–traM* | Δ28,988 bp (x5) | 30 genes in transfer region | B | GTGGCCGCCG |
| | | | *trbE–kfrA* | Δ30,679 bp (x6) | 33 genes in transfer and maintenance/control regions | C | |
| | | $t_{28}$ | *trbE–traM* [3] | Δ28,549 bp (x6) | 30 genes in transfer region | A | GGCCAaCTC* |
| | | | *kfrA* [2] | Δ84 bp | KfrA protein in maintenance/control region | A | |
| | | | *trbE–traM* | Δ28,988 bp (x6) | 30 genes in transfer region | B | GTGGCCGCCG |
| | | | *trbE–kfrA* | Δ30,679 bp (x6) | 33 genes in transfer and maintenance/control regions | C | |
| **Tet-** | Starting point | $t_0$ | *klcB* | Δ144 bp | KlcB protein in maintenance/control region | A | |
| | Biofilm | $t_{14}$ | *trbL* | Δ219 bp | putative DNA topoisomerase in transfer region | A | |
| | | | *trbL* | Δ306 bp | putative DNA topoisomerase | B | |
| | | | *trbB* | SNP | conjugal transfer protein TrbB in transfer region | B | |
| | | | *trbI* | Δ648 bp | conjugal transfer protein TrbI in transfer region | B | |
| | | | *trbL–trbN* | Δ1,881 bp (x2) | 3 genes in transfer region | A | |
| | | | *trbB–traM* | Δ31,451 bp | 33 genes in transfer region | C | CGCGGCGC |
| | | | *trbB–traC* | Δ21,755 bp | 23 genes in transfer region | C | CCTGGTCGGCGA |
| | Planktonic | $t_6$ [4] | *trbE–incC2* | Δ34,594 bp (x4) | 36 genes in transfer and maintenance/control regions | B | GGCCGCCTGG |
| | | $t_{10}$ [4] | *trbA–traD* | Δ23,369 bp (x4) | 25 genes in transfer region | C | |
| | | | *trbE–kfrA* | Δ31,221 bp (x2) | 33 genes in in transfer and maintenance/control regions | C | |
| | | $t_{14}$ | *trbC–kfrA* | Δ32,964 bp(x6) | 35 genes in transfer and maintenance/control regions | A | CGGTGGCCTTC |

[1] The numbers in parentheses (x2, x4, x5, x6, x7) refer to the number of clones that showed that exact genetic change. When two genetic changes are shown for the same plasmid gene or region at a particular time point, these were observed in separate clones.

[2] These mutations were found in one clone that also has the mutation listed right above

[3] A plasmid with this genotype was tested in other bacterial species (Fig. 5).

[4] Plasmid-containing clones were no longer detected after Day 6 (or 10), explaining why clones were sequenced at these time points in planktonic populations without tetracycline

* Direct repeat right had a 'G' instead of an 'A'

Supplementary Table 2: Matings with various *A. baumanni* clones as donor and *E. coli* as recipient show that loss of transfer genes disabled conjugative plasmid transfer

| Donor | Recipient | D (cfu/ml) [a] | R (cfu/ml) [a] | T (cfu/ml) [a] | T/R | Standard deviation | 95% CI | T/D | Standard deviation | 95% CI |
|---|---|---|---|---|---|---|---|---|---|---|
| Ancestor | *E. coli* K12Nal[b] | 2.7E+09 | 6.8E+07 | 3.4E+03 | 5.3E-05 | 2.9E-05 | 7.3E-05 | 1.3E-06 | 6.3E-07 | 1.6E-06 |
| Evolved clone with intact plasmid | " " | 9.7E+08 | 6.5E+07 | 3.4E+02 | 5.4E-06 | 1.7E-06 | 4.2E-06 | 3.6E-07 | 1.2E-07 | 2.9E-07 |
| Evolved clone with truncated plasmid | " " | 2.0E+09 | 6.4E+07 | <10 | <1.6E-07 | - | - | - | - | - |

[a] Values are averages from three replicate mating assays

[b] E. coli K12 Nal is a nalidixic acid resitant mutant of MG1655

Supplementary Table 3: List of all sequenced strains with their predicted mutations generated by Breseq using the ‘gdtools COMPARE’ subcommand.

**Predicted mutations**

| seq id | position | mutation | DOD1508 | DOD1509 | DOD1510 | DOD1511 | DOD1512 | DOD408 | DOD409 | DOD410 | DOD-67 | annotation | gene | description |
|---|---|---|---|---|---|---|---|---|---|---|---|---|---|---|
| AJ564903 | 340 | Δ23,369 bp | | | | | | | | | | | *[trbA]–[traD]* | **25 genes** Show |
| AJ564903 | 566 | Δ31,764 bp | | | | | | | | | | | *trbB–[traL]* | **32 genes** Show |
| AJ564903 | 771 | G→A | | | | | | | | | | M1M (ATG→ATA) † | *trbB* → | putative ATPase, autophosphorylase |
| AJ564903 | 904 | Δ32,730 bp | | | | | | | | | | | *[trbB]–[traN]* | **34 genes** Show |
| AJ564903 | 1,452 | Δ21,755 bp | | | | | | | | | | | *[trbB]–[traC]* | **23 genes** Show |
| AJ564903 | 1,669 | Δ31,451 bp | | | | | | | | | | | *[trbB]–[traM]* | **33 genes** Show |
| AJ564903 | 1,937 | Δ32,964 bp | | | | | | | | | | | *[trbC]–[kfrA]* | **35 genes** Show |
| AJ564903 | 2,309 | Δ33,164 bp | | | | | | | | | | | *[trbD]–[kfrA]* | **34 genes** Show |
| AJ564903 | 2,731 | Δ34,594 bp | | | | | | | | | | | *[trbE]–[incC1]* | **36 genes** Show |
| AJ564903 | 3,700 | Δ30,220 bp | | | | | | | | | | | *[trbE]–[traN]* | **31 genes** Show |
| AJ564903 | 3,704 | Δ31,221 bp | | | | | | | | | | | *[trbE]–[kfrA]* | **33 genes** Show |
| AJ564903 | 4,181 | Δ28,988 bp | | | | | | | | | | | *[trbE]–[traM]* | **30 genes** Show |
| AJ564903 | 4,301 | Δ30,679 bp | | | | | | | ? | 100% | | | *[trbE]–[kfrA]* | **33 genes** Show |
| AJ564903 | 4,731 | Δ28,549 bp | | | | | | | ? | Δ | | | *[trbE]–traM* | **30 genes** Show |
| AJ564903 | 5,481 | Δ29,446 bp | | | | | | | ? | Δ | | | *[trbF]–[kfrA]* | **32 genes** Show |
| AJ564903 | 7,563 | Δ648 bp | | | | | | ? | ? | Δ | | coding (290-937/1422 nt) | *trbI* → | TrbI protein |
| AJ564903 | 9,268 | $(GCGCAGGGC)_{2\to1}$ | | | | | | ? | ? | Δ | | coding (553-561/765 nt) | *trbJ* → | TrbJ protein |
| AJ564903 | 10,666 | Δ219 bp | | | | | | ? | ? | Δ | | coding (939-1157/1719 nt) | *trbL* → | putative DNA topoisomerase |
| AJ564903 | 10,697 | Δ1,881 bp | | | | | | ? | ? | Δ | | | *[trbL]–[trbN]* | *[trbL]*, *trbM*, *[trbN]* |
| AJ564903 | 10,972 | Δ405 bp | | | | | | ? | ? | Δ | | coding (1245-1649/1719 nt) | *trbL* → | putative DNA topoisomerase |
| AJ564903 | 35,210 | Δ84 bp | | | | | | | | | | coding (336-419/1032 nt) | *kfrA* ← | KfrA protein |
| AJ564903 | 35,211 | Δ17 bp | | | | | | | | | | coding (402-418/1032 nt) | *kfrA* ← | KfrA protein |
| AJ564903 | 35,225 | Δ141 bp | | | | | | | | | | coding (264-404/1032 nt) | *kfrA* ← | KfrA protein |
| AJ564903 | 35,241 | G→A | | | | | | | | | | L130L (CTG→TTG) | *kfrA* ← | KfrA protein |
| AJ564903 | 35,338 | G→C | | | | | | | | | | A97A (GCC→GCG) | *kfrA* ← | KfrA protein |
| AJ564903 | 40,050 | Δ144 bp | | | | 100% | | | | | | coding (579-722/1209 nt) | *klcB* ← | KlcB protein |
| AJ564903 | 46,936 | A→G | | | | | | | | | | pseudogene (10638/14051 nt) pseudogene (5865/8895 nt) | *tnpA* ← *tnpA* ← | hypothetical protein hypothetical protein |
| CP000521 | 17,639 | $(TTCTTTGGA)_{11\to9}$ | | | | | | | ? | | | intergenic (+247/+1487) | *A1S_0014* → / ← *A1S_3477* | tyrosyl-tRNA synthetase/hypothetical protein |
| CP000521 | 17,648 | $(TTCTTTGGA)_{11\to10}$ | | | | | | | ? | | | intergenic (+256/+1487) | *A1S_0014* → / ← *A1S_3477* | tyrosyl-tRNA synthetase/hypothetical protein |
| CP000521 | 17,656 | $(TTCTTTGGA)_{11\to12}$ | | | | | | | | | | intergenic (+264/+1487) | *A1S_0014* → / ← *A1S_3477* | tyrosyl-tRNA synthetase/hypothetical protein |
| CP000521 | 17,656 | $(TTCTTTGGA)_{11\to13}$ | | | | | | | | | | intergenic (+264/+1487) | *A1S_0014* → / ← *A1S_3477* | tyrosyl-tRNA synthetase/hypothetical protein |
| CP000521 | 17,656 | $(TTCTTTGGA)_{11\to16}$ | | | | | | | | | | intergenic (+264/+1487) | *A1S_0014* → / ← *A1S_3477* | tyrosyl-tRNA synthetase/hypothetical protein |
| CP000521 | 17,656 | $(TTCTTTGGA)_{11\to17}$ | | | | | | | | | | intergenic (+264/+1487) | *A1S_0014* → / ← *A1S_3477* | tyrosyl-tRNA synthetase/hypothetical protein |
| CP000521 | 17,656 | $(TTCTTTGGA)_{11\to18}$ | 100% | | 100% | 100% | | | | 100% | | intergenic (+264/+1487) | *A1S_0014* → / ← *A1S_3477* | tyrosyl-tRNA synthetase/hypothetical protein |
| CP000521 | 17,656 | $(TTCTTTGGA)_{11\to19}$ | | | | | | | | | | intergenic (+264/+1487) | *A1S_0014* → / ← *A1S_3477* | tyrosyl-tRNA synthetase/hypothetical protein |
| CP000521 | 101,403 | A→G | 100% | | | 100% | 100% | 100% | 100% | 100% | | intergenic (-614/-2418) | *A1S_0081* ← / → *A1S_0086* | putative signal peptide/hypothetical protein |
| CP000521 | 101,415 | T→C | | | | 100% | | 100% | 100% | 100% | | intergenic (-626/-2406) | *A1S_0081* ← / → *A1S_0086* | putative signal peptide/hypothetical protein |
| CP000521 | 149,238 | Δ122 bp | | | | | | | | | | | *[trnG]* | *[trnG]* |
| CP000521 | 485,616 | C→T | | | | | | | | | | A375T (GCA→ACA) | *A1S_0446* ← | putative transport protein (CPA2 family) |
| CP000521 | 486,124 | C→T | | | | | | | | | | M205I (ATG→ATA) | *A1S_0446* ← | putative transport protein (CPA2 family) |
| CP000521 | 511,070 | $(ATTTTGAAC)_{2\to3}$ | | | | | | | | | | coding (168/1698 nt) | *A1S_0472* ← | 2-isopropylmalate synthase |
| CP000521 | 516,966 | Δ4 bp | | | | | | | | | | coding (252-255/606 nt) | *A1S_0476* → | ATP-dependent Clp protease proteolytic subunit |
| CP000521 | 517,105 | Δ1 bp | | | | | | 100% | 100% | 100% | | coding (391/606 nt) | *A1S_0476* → | ATP-dependent Clp protease proteolytic subunit |
| CP000521 | 826,432 | Δ19 bp | | ? | ? | | ? | 100% | | | | coding (315-333/1227 nt) | *A1S_0694* → | FilE |
| CP000521 | 826,432 | Δ10 bp | | ? | ? | | ? | Δ | | | | coding (315-324/1227 nt) | *A1S_0694* → | FilE |
| CP000521 | 826,432 | Δ1 bp | | ? | ? | | ? | Δ | ? | | | coding (315/1227 nt) | *A1S_0694* → | FilE |
| CP000521 | 826,440 | Δ2 bp | | ? | ? | | ? | Δ | | | | coding (323-324/1227 nt) | *A1S_0694* → | FilE |
| CP000521 | 826,441 | Δ1 bp | | ? | ? | | ? | Δ | | | | coding (324/1227 nt) | *A1S_0694* → | FilE |
| CP000521 | 826,444 | Δ3 bp | | ? | ? | | ? | Δ | | | | coding (327-329/1227 nt) | *A1S_0694* → | FilE |
| CP000521 | 826,448 | Δ2 bp | | ? | ? | | ? | Δ | | | | coding (331-332/1227 nt) | *A1S_0694* → | FilE |
| CP000521 | 877,534 | 2 bp→CA | | | | | | | | | | coding (2165-2166/2940 nt) | *A1S_0742* → | iron-regulated protein |
| CP000521 | 877,535 | T→A | | | | | | | | | | I722I (ATT→ATA) | *A1S_0742* → | iron-regulated protein |
| CP000521 | 877,895 | G→A | | | | | | | | | | L842L (CTG→CTA) | *A1S_0742* → | iron-regulated protein |
| CP000521 | 877,958 | A→T | | | | | | | | 100% | | G863G (GGA→GGT) | *A1S_0742* → | iron-regulated protein |
| CP000521 | 880,449 | C→T | | | | | | | | | | G575D (GGC→GAC) | *A1S_0745* ← | hypothetical protein |
| CP000521 | 1,029,854 | $(TGGCGA)_{2\to3}$ | | | | | | 100% | 100% | 100% | | coding (743/1419 nt) | *A1S_0887* → | phosphomannomutase |
| CP000521 | 1,030,421 | T→C | | | | | | | | | | V437A (GTA→GCA) | *A1S_0887* → | phosphomannomutase |
| CP000521 | 1,075,007 | T→G | | | | | | | | | | intergenic (-102/+7) | *A1S_0924* ← / ← *A1S_0925* | choline dehydrogenase/choline dehydrogenase |
| CP000521 | 1,075,007 | 2 bp→GT | 100% | 100% | 100% | 100% | 100% | 100% | 100% | 100% | | intergenic (-102/+6) | *A1S_0924* ← / ← *A1S_0925* | choline dehydrogenase/choline dehydrogenase |
| CP000521 | 1,075,008 | G→T | | | | | | | | | | intergenic (-103/+6) | *A1S_0924* ← / ← *A1S_0925* | choline dehydrogenase/choline dehydrogenase |
| CP000521 | 1,075,010 | C→G | | | | | | | | | | intergenic (-105/+4) | *A1S_0924* ← / ← *A1S_0925* | choline dehydrogenase/choline dehydrogenase |
| CP000521 | 1,075,010 | 2 bp→GC | 100% | 100% | 100% | 100% | 100% | 100% | 100% | 100% | | intergenic (-105/+3) | *A1S_0924* ← / ← *A1S_0925* | choline dehydrogenase/choline dehydrogenase |
| CP000521 | 1,075,011 | G→C | | | | | | | | | | intergenic (-106/+3) | *A1S_0924* ← / ← *A1S_0925* | choline dehydrogenase/choline dehydrogenase |
| CP000521 | 1,217,951 | C→T | | | | | | | | | | Q269* (CAA→TAA) | *A1S_1055* → | hypothetical protein |
| CP000521 | 1,318,577 | G→T | | | | | | | | | | V96V (GTC→GTA) | *A1S_1127* ← | oxidoreductase short chain dehydrogenase/reductase family |
| CP000521 | 1,353,821 | G→A | | | | | | | | | | G84S (GGT→AGT) | *A1S_1158* → | putative signal peptide |
| CP000521 | 1,386,215 | $(T)_{8\to9}$ | | | | | | | | | | intergenic (-499/-12) | *A1S_1182* ← / → *A1S_1183* | cyclic AMP receptor protein/hypothetical protein |
| CP000521 | 1,630,443 | C→T | | | | ? | | | 100% | ? | | pseudogene (378/706 nt) | *A1S_3662* → | hypothetical protein |
| CP000521 | 1,762,595 | T→C | | | | | | | | | | D174D (GAT→GAC) | *A1S_1513* → | putative membrane protein |
| CP000521 | 1,885,761 | C→T | | | | | | | | | | T232I (ACT→ATT) | *A1S_1616* → | hypothetical protein |
| CP000521 | 2,193,897 | G→A | | 100% | | | | | ? | ? | | D173D (GAC→GAT) | *A1S_1893* ← | 3-oxoadipate CoA-transferase subunit B |
| CP000521 | 2,212,578 | C→T | | | | | | | | | | G405D (GGT→GAT) | *A1S_1910* ← | ATP-binding protease component |
| CP000521 | 2,213,307 | G→T | | | | | | | | | | S162* (TCA→TAA) | *A1S_1910* ← | ATP-binding protease component |
| CP000521 | 2,292,757 | +122 bp | | | | | | | | | | intergenic (+265/-11) | *A1S_1978* → / → *A1S_1979* | response regulator protein/putative transcriptional regulator |
| CP000521 | 2,374,353 | A→G | | | | | | | | | | intergenic (+94/-109) | *A1S_3775* → / → *A1S_2038* | hypothetical protein/hypothetical protein |
| CP000521 | 2,396,955 | $(GGTTCA)_{17\to19}$ | | | | | | | | | | coding (1230/2109 nt) | *A1S_2056* ← | DNA polymerase III tau and gamma subunits (DNA e longation factor III) |
| CP000521 | 2,422,982 | C→T | | | | | | | | | | intergenic (-34/+290) | *A1S_2076* ← / ← *A1S_2077* | putative outer membrane porin receptor for Fe(II I)-coprogen, Fe(III)-ferrioxamine B and Fe(III)-rhodotrul i c acid uptake (fhuE)/putative outer membrane porin receptor for Fe(II I)-coprogen, Fe(III)-ferrioxamine B and Fe(III)-rhodotrul i c acid uptake (fhuE) |
| CP000521 | 2,466,306 | G→T | | | | | | | | | | Q41K (CAA→AAA) | *A1S_3785* ← | hypothetical protein |
| CP000521 | 2,576,310 | T→C | | | | | | | | | | E175G (GAG→GGG) | *A1S_2213* ← | CsuE |
| CP000521 | 2,576,373 | A→G | | | | | | | | | | L154S (TTA→TCA) | *A1S_2213* ← | CsuE |
| CP000521 | 2,576,530 | T→C | | | | | | | | | | S102G (AGT→GGT) | *A1S_2213* ← | CsuE |
| CP000521 | 2,576,533 | C→A | | | | | | | | | | D101Y (GAC→TAC) | *A1S_2213* ← | CsuE |
| CP000521 | 2,576,538 | A→G | | | | | | | | | | I99T (ATT→ACT) | *A1S_2213* ← | CsuE |
| CP000521 | 2,576,586 | G→A | | | | | | | | | | S83L (TCG→TTG) | *A1S_2213* ← | CsuE |
| CP000521 | 2,576,605 | A→T | | | | | | | | | | Y77N (TAT→AAT) | *A1S_2213* ← | CsuE |
| CP000521 | 2,708,492 | T→C | | | | | | | | | | L817S (TTA→TCA) | *A1S_2337* → | sensory box protein |
| CP000521 | 2,708,717 | T→A | | | | | | | | | | M892K (ATG→AAG) | *A1S_2337* → | sensory box protein |
| CP000521 | 2,708,737 | A→G | | | | | | | | | | M899V (ATG→GTG) | *A1S_2337* → | sensory box protein |
| CP000521 | 3,059,254 | G→A | | | | | | | | | | Q168* (CAA→TAA) | *A1S_2642* ← | regulatory protein TetR |
| CP000521 | 3,059,463 | C→A | | | | | | | | | | C98F (TGT→TTT) | *A1S_2642* ← | regulatory protein TetR |
| CP000521 | 3,093,102 | G→A | | | | | | | | | | pseudogene (398/1836 nt) | *A1S_2668* → | phosphoenolpyruvate carboxykinase |
| CP000521 | 3,093,890 | G→A | | | | | | | | | | pseudogene (1186/1836 nt) | *A1S_2668* → | phosphoenolpyruvate carboxykinase |
| CP000521 | 3,129,237 | A→G | | | | | | | ? | ? | | G15G (GGT→GGC) | *A1S_2696* ← | hypothetical protein |
| CP000521 | 3,306,773 | $(T)_{8\to7}$ | | | | | | 100% | 100% | 100% | | coding (304/1212 nt) | *A1S_2858* ← | putative methyltransferase |
| CP000521 | 3,328,022 | T→C | | | | | | | | | | V168A (GTC→GCC) | *A1S_2878* → | hypothetical protein |
| CP000521 | 3,434,070 | Δ2 bp | | | | | | | | | | coding (886-887/1194 nt)\|coding (4-5/105 nt) | *A1S_3863* → *A1S_3862* ← | hypothetical protein hypothetical protein |
| CP000521 | 3,441,396 | Δ1 bp | 100% | | | 100% | | 100% | 100% | 100% | | intergenic (-28/+371) | *A1S_r12* ← / ← *A1S_2955* | 16S ribosomal RNA/peptidyl prolyl-cis-trans-isomerase rotamase |
| CP000521 | 3,468,548 | $(ATGGTG)_{9\to10}$ | | | | | | | | | | coding (454/987 nt) coding (193/231 nt) | *A1S_2977* ← *A1S_3873* → | cation diffusion facilitator family transporter hypothetical protein |
| CP000521 | 3,468,548 | $(ATGGTG)_{9\to11}$ | | | | | | | | | | coding (454/987 nt) coding (193/231 nt) | *A1S_2977* ← *A1S_3873* → | cation diffusion facilitator family transporter hypothetical protein |
| CP000521 | 3,500,412 | T→C | | | | | | | | | | intergenic (-119/+6) | *A1S_3014* ← / ← *A1S_r13* | hypothetical protein/5S ribosomal RNA |
| CP000521 | 3,500,576 | G→A | | | | | | | | | | intergenic (-25/+140) | *A1S_r13* ← / ← *A1S_r14* | 5S ribosomal RNA/23S ribosomal RNA |
| CP000521 | 3,500,617 | Δ1 bp | | | | | | | | | | intergenic (-66/+99) | *A1S_r13* ← / ← *A1S_r14* | 5S ribosomal RNA/23S ribosomal RNA |
| CP000521 | 3,500,634 | C→T | | | | | | | | | | intergenic (-83/+82) | *A1S_r13* ← / ← *A1S_r14* | 5S ribosomal RNA/23S ribosomal RNA |
| CP000521 | 3,870,029 | Δ21 bp | | | | | | | | | | pseudogene (577-597/605 nt) | *A1S_3363* ← | membrane metalloendopeptidase protein |
| CP000521 | 3,870,072 | $(CTCCAGATG)_{5\to6}$ | | | | | | | | | | pseudogene (554/605 nt) | *A1S_3363* ← | membrane metalloendopeptidase protein |
| CP000521 | 3,870,072 | $(CTCCAGATG)_{5\to13}$ | | | | | | | | | | pseudogene (554/605 nt) | *A1S_3363* ← | membrane metalloendopeptidase protein |
| CP000521 | 3,928,225 | $(G)_{7\to6}$ | | | | | | | | | | coding (28/780 nt) | *A1S_3417* → | regulatory proteins IclR |
| CP000521 | 3,928,225 | $(G)_{7\to8}$ | | | | | | | | | | coding (28/780 nt) | *A1S_3417* → | regulatory proteins IclR |
| CP000521 | 3,962,192 | T→C | | | | | | | | | | L393P (CTG→CCG) | *A1S_3449* → | phosphoenolpyruvate carboxylase |
